# plASgraph - using graph neural networks to detect plasmid contigs from an assembly graph

**DOI:** 10.1101/2022.05.24.493339

**Authors:** Janik Sielemann, Katharina Sielemann, Broňa Brejová, Tomáš Vinař, Cedric Chauve

**Affiliations:** Computational Biology, Faculty of Biology, Center for Biotechnology (CeBiTec) & Graduate School, DILS, Bielefeld Institute for Bioinformatics Infrastructure (BIBI), Bielefeld University, 33615 Bielefeld, Germany; Department of Mathematics, Simon Fraser University, Burnaby, Canada; Genetics and Genomics of Plants, Faculty of Biology, Center for Biotechnology (CeBiTec) & Graduate School DILS, Bielefeld Institute for Bioinformatics Infrastructure (BIBI), Bielefeld, University, 33615 Bielefeld, Germany; Department of Computer Science, Faculty of Mathematics, Physics and Informatics, Comenius University in Bratislava, Slovakia; Department of Applied Informatics, Faculty of Mathematics, Physics and Informatics, Comenius University in Bratislava, Slovakia

**Keywords:** Contig classification, graph neural network, machine learning, plasmids

## Abstract

Identification of plasmids from sequencing data is an important and challenging problem related to antimicrobial resistance spread and other One-Health issues. In our work, we provide a new architecture for identifying plasmid contigs in fragmented genome assemblies built from short-read data. Unlike previous machine-learning approaches for this problem, which classify individual contigs separately, we employ graph neural networks (GNNs) to include information from the assembly graph. Propagation of information from nearby nodes in the graph allows accurate classification of even short contigs that are difficult to classify based on sequence features or database searches alone.

Our new species-agnostic software tool plASgraph outperforms recently developed PlasForest, which uses database searches to supplement sequence-based features. Since our tool does not rely on existing plasmid databases, it is more suitable for classification of contigs in novel species and discovery of previously unknown plasmid sequences. Our tool can also be trained on a specific species, and in that scenario it outperforms mlplasmids trained on the same species.

On one hand, our work provides a new, accurate, and easy to use tool for plasmid classification; on the other hand, it serves as a motivation for more widespread use of GNNs in bioinformatics, such as in pangenome sequence analysis, where sequence graphs serve as a fundamental data structure.

**Availability:** https://github.com/cchauve/plASgraph

## 1 Introduction

Plasmids are mobile genetic elements that are involved in horizontal gene transfers and have been shown to be a major vector for the spread of antimicrobial resistance (AMR) genes [8, 20]. Plasmids are extra-chromosomal DNA molecules, often circular and significantly shorter than bacterial chromosomes, and can occur in multiple copies in a bacterial cell. Whereas some bacteria do not contain any plasmid, it is common to observe several plasmids co-existing within a bacterial cell, often with different copy numbers. Due to their high mobility and impact in AMR spread, the detection of plasmids from sequencing data is an important question in One-Health epidemiologic surveillance approaches, see e.g. [10].

Given sequencing data, either from a bacterial isolate or from a metagenome, the detection of plasmids can be approached at various levels of detail. The most elementary task, *contig classification*, aims at detecting which assembled contigs are likely to originate from a plasmid. *Plasmid binning* aims at grouping contigs into groups likely to originate from the same plasmid. Last, *plasmid assembly* aims at reconstructing full plasmid sequences. While obtaining full plasmids provides the most accurate information, the ability to extract plasmid contigs from assembled sequencing data (the contig classification problem) already provides very useful information, allowing e.g. to identify genes that might be susceptible to transfer to other bacteria. Moreover, the prediction of plasmid contigs can be used as an input for plasmid binning or assembly. For example, the plasmid binning method gplas [4] relies on a preliminary contig classification obtained with mlplasmids [5] and the metagenome plasmid assembly method SCAPP [22] relies on classifying contigs using PlasClass [21].

While the analysis of plasmids from sequencing data has been a very active research area, the problems mentioned above are still challenging, especially when sequencing data are provided in the form of Illumina short reads [6]. In the present paper, we propose a novel method for the contig classification problem, specifically designed to analyse short-read contigs from a single bacterial isolate.

There exists a large corpus of algorithms for the contig classification problem, most of them developed recently. These methods rely mainly on machine-learning approaches. The earliest method for contig classification was cBar [30], which introduced the use of the *k*-mer profile of a contig as the main feature in a machine-learning classification model; in cBar, the model was trained on a large dataset of closed bacterial genome assemblies. The general principle of using *k*-mer properties as classification features has also been used in several recent machine-learning classifiers, namely PlasFlow [16], mlplasmids [5], and PlasClass [21]. PPR-Meta [11] is a deep-learning method that relies on one-hot encoded contig sequences. PlasForest [23] and Deeplasmid [3] are two recent methods based on machine-learning models that use different features for a given contig, such as its GC content (generally plasmids have a GC content different from chromosomes) and the presence of plasmid-specific sequences, detected through the mapping against a reference plasmid database. RFPlasmid [27] combines both kinds of features, the *k*-mer profile and plasmid-specific sequences. Among the methods introduced above, both mlplasmids and RFPlasmid are species-specific methods, i.e. require a model to be trained per bacterial species; in contrast, PlasFlow, PlasClass, PlasForest and Deeplasmid are tools that do not target a specific species.

The recent method 3CAC [24] introduced the idea that the classification of a contig can be improved from the knowledge of the classification of the neighbouring contigs in the assembly graph. Most current assembly programs [7, 28] output an assembly graph containing final contigs as nodes and possible connections between them supported by sequencing data as edges. Individual molecules, such as chromosomes or plasmids, ideally correspond to walks in this graph, but some edges may be missing, disconnecting the walk. Conversely, the walks for individual molecules often form complicated tangled structures joined at shared and repeated sequences. Nonetheless, adjacent nodes often share the same molecule of origin and thus the same class. 3CAC applies simple heuristics to improve machine learning predictions for individual contigs based on their adjacency in the graph. Our aim is to integrate the information from the assembly graph directly to the underlying machine learning model.

Here, we introduce a novel machine-learning method, plASgraph, for the problem of classifying short-read contigs as plasmidic or chromosomal. Our method is based on combining features of existing methods with a novel approach incorporating a graph neural network (GNN) [12]. More precisely, plASgraph associates to each contig of a bacterial genome assembly a set of features that have been shown to differentiate plasmids and chromosomes: read coverage, used as a proxy of copy number, GC content and contig length, together with two novel features, the node degree in the assembly graph and the distance between the contig *k*-mer profile and the whole assembly *k*-mer profile. The rationale to integrate the *k*-mer profile by comparing it to the assembly-wide profile is to allow our model to be species-agnostic, i.e. not learning a species-specific *k*-mer profile, as is done in species-specific models such as mlplasmids and RFPlasmid. Moreover, plASgraph is a *de novo* tool that does not require the comparison of the input contigs with a database of known plasmids. Based on these features, plASgraph trains a GNN model whose core is a set of graph convolutional layers aimed at propagating the information from neighbouring contigs in the assembly graph. To the best of our knowledge, plASgraph is the first method that applies GNNs to contig classification in an assembly graph, building on the idea (introduced in 3CAC) that information from neighbouring contigs can improve accuracy. Outside of classification, GNNs were also used recently on assembly graphs for metagenomic contigs binning [17].

The output of plASgraph is a pair of scores for each graph node, a plasmid score and a chromosomal score, used to determine if a given contig is likely to originate from a plasmid or a chromosome or both. Unlike other methods, the two scores associated to a contig allow to detect *ambiguous* contigs that have shared sequences of both plasmidic and chromosomal origin. In order to train plASgraph, we rely on the availability of hybrid sequencing data composed of short and long reads; our method thus does not depend on the availability of a reference plasmid database.

We evaluated the performance of plASgraph in two contexts: species-specific and species-agnostic. In the species-specific context, we trained plASgraph on data from a single bacterial species and compared the model accuracy to mlplasmids [5] when applied to isolates from the same species and from other species. In the species-agnostic context, we trained plASgraph on data combined from several species and compared it to PlasForest [23], which in addition to the sequence-based features also uses the information from database searches. In both scenarios, plASgraph outperforms the competing tool.

## 2 Methods

### 2.1 Overview

The input to our problem is an assembly graph of a bacterial isolate in which nodes correspond to contigs and edges correspond to adjacencies supported by sequencing data. This graph is typically created from short reads and as a result can contain a large number of contigs of various sizes. For example, in our *E. faecium* training set (average genome size 2.84 Mbp), the number of contigs ranged from tens to hundreds with an N50 value between 34 kbp and 253 kbp. Our goal is to classify individual contigs as originating from a plasmid or from a chromosome. However, some contigs in fact correspond to sequences that occur as parts of both plasmids and chromosomes within the same sample (for example, mobile elements or low-complexity sequences); we call such contigs *ambiguous*. Due to the presence of ambiguous contigs, we treat the problem as two separate classification tasks, generating independent scores for chromosome and plasmid labels.

Ideally, the input assembly graph would consist of a few connected components, each corresponding either to a single chromosome or a plasmid. In a fully resolved assembly, each chromosome and plasmid would actually correspond to an isolated vertex. However, in graphs created from short-read data, walks corresponding to individual molecules are typically interconnected by spurious edges or through ambiguous contigs, creating complex structures (see Figure 5 for an illustration). The main novelty of our approach is to use the graph neighbourhood of a node as a source of information, under the assumption that other contigs from the same molecule (chromosome or plasmid) are likely to belong to this neighbourhood.

Given a comprehensive database of closed genomes for a given bacterial species, longer contigs can likely be classified simply based sequence homology; in our work, we concentrate on classifying contigs in a *de novo* framework that does not rely on existing reference genomes. The *de novo* contigs classification problem is of interest for e.g. the analysis of samples from poorly sampled bacterial species.

### 2.2 Input features

As an input to the classification task, each contig is characterized by five input features:

1. the *degree* of the corresponding node in the assembly graph;
2. the *relative contig length*, defined by the contig length divided by the length of the longest contig;
3. the *relative GC content*, defined by subtracting the average GC content (expressed as a percentage) of the whole assembly from the contig GC content;
4. the *relative coverage*, defined as the contig read depth divided by the median read depth over the whole assembly (in our experiments, we use the read depth value provided by the Unicycler assembler [28]);
5. the *relative pentamer distance*, defined as the Euclidean distance between the pentamer profile of the contig and the pentamer profile of the whole assembly; we define the pentamer profile of a contig or a set of contigs as the count vector for all pentamers (reverse complements were aggregated), shifted and scaled so that the smallest vector entry is 0 and the largest vector entry is 1.

The motivation to rely on relative features instead of absolute features is to enable the model to generalize across species, and thus to not be dependent on species-specific values. Regarding the relative pentamer distance, one can expect that large chromosomal contigs will have a value closer to zero, while shorter plasmid contigs will exhibit a large value for this feature. Moreover, by abstracting the pentamer content of a sample by the relative pentamer distance, we expect that our model will be less susceptible to learning to classify chromosome sequences by simply recognizing the pentamer frequencies characteristic for a particular species or a clade.

### 2.3 Model architecture

We employed a deep neural network to solve our classification task. The key part of our architecture is the use of a *graph convolutional network* (GNN) [15] to account for the assembly graph structure. The propagation of information between individual nodes is accomplished by *graph convolutional layers* (GCLs). Briefly, the input to a GCL contains a vector of *k* features for each of the *n* nodes of the graph and the adjacency matrix of the graph. The layer first combines the feature vectors corresponding to the node and its neighbours, with weight of nodes depending on their degree. The feature vector of each node is then transformed by a fully-connected layer with *ℓ* output features followed by a non-linear activation. More precisely, if we organize the *n* feature vectors into the *n* × *k* matrix *X*, the graph convolutional layer can be expressed as

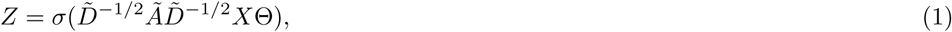

where *Ã* is the graph adjacency matrix with ones along the diagonal, 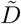 is a diagonal matrix where 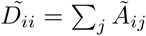, Θ is a *k* × *ℓ* matrix of trainable weights, *σ* is a non-linear activation function, and *Z* is the *n* × *ℓ* matrix of output feature vectors. A single GCL integrates information from an immediate neighbourhood of a node; by employing *d* GCLs one integrates the information from the distance of at most *d* for each node.

Figure 1 shows the architecture we have designed for our task. The five input features for each node are first transformed by a fully connected layer to a vector of length 32 per node. This is followed by six GCLs using the same weight matrix Θ. The last two fully connected layers operate on each node separately, the first producing a vector of length 32, and the second producing two output scores, loosely interpretable as probabilities of the node being part of a chromosome and plasmid, respectively. Since these two outputs correspond to two separate classification tasks, we do not require these two scores to sum to one. Each layer is followed by the ReLU activation, except for the last layer, which uses the sigmoid activation. All layers excluding the last one are followed by 10% dropout to prevent overfitting.

**Figure 1.**
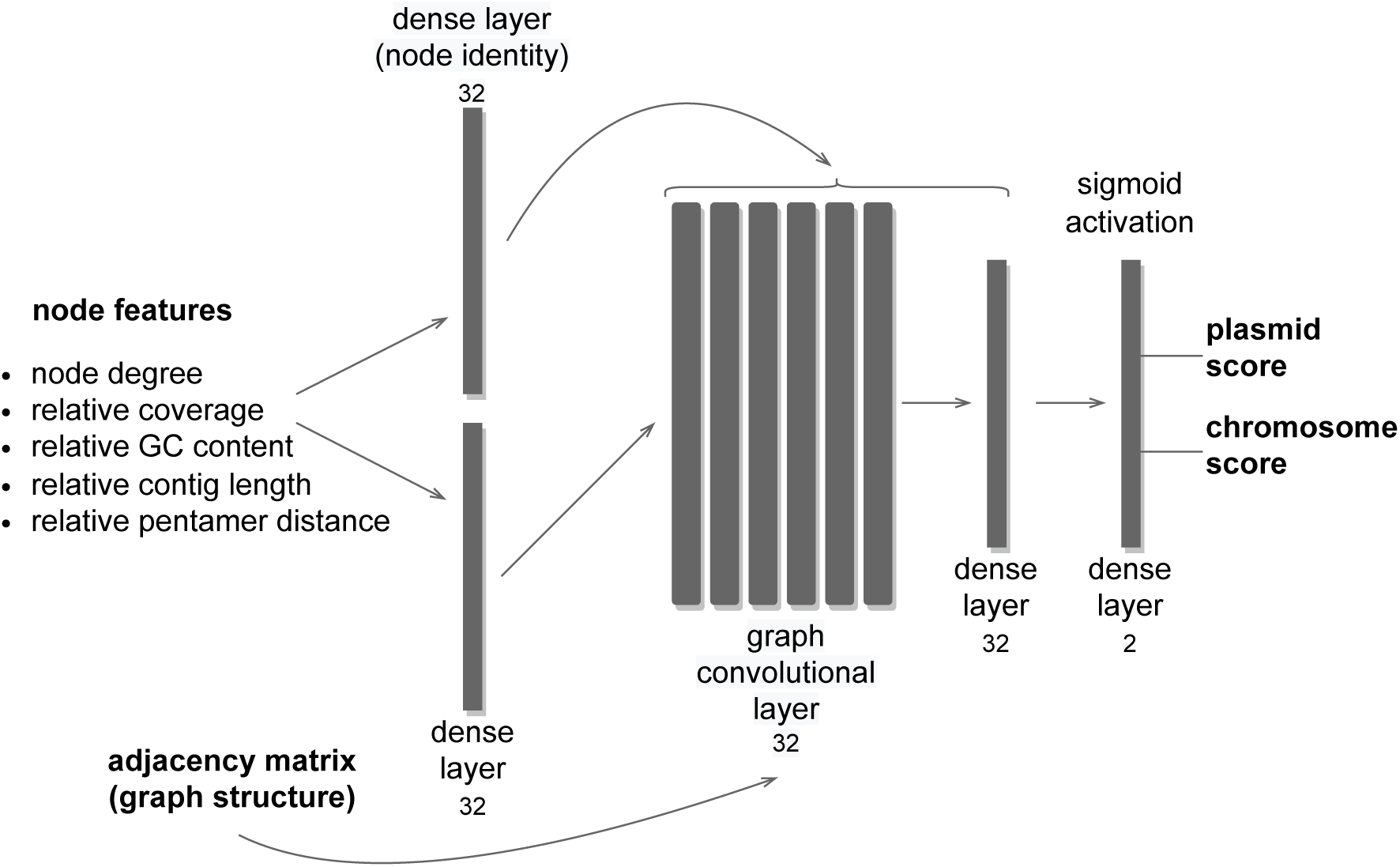
Model architecture of plASgraph. The model uses as inputs the assembly graph structure and five input features per node (contig). The core of the architecture is composed of six graph convolutional layers. The model generates two outputs per node, which facilitate the classification of plasmids and chromosomes as two separate classification tasks.

GCLs combine features of each node with features of the neighbours and over time, the influence of the original features is greatly diminished. In our task, the original features can be highly informative, especially for nodes corresponding to longer contigs; therefore we want to maintain node identity throughout the computation. To accomplish this, each GCL and the penultimate dense layer receive as an input an additional vector of length 32 for each node, representing a separate encoding of the five input features. Thus, each of these layers starts with the input vectors of length 64 and reduces them to vectors of length 32.

The network is trained using Adam optimizer [14] with binary cross entropy loss function and a constant learning rate of 0.005. The model is implemented using Keras [9] and TensorFlow v2.8.0 [1], with GCLs from Spektral v1.0.8 [12].

### 2.4 Data preparation

For training and testing, we used data from eight bacterial species, listed in Tab. 1. Individual bacterial isolates were sequenced both by short-read (Illumina) and long-read (Oxford Nanopore or Pacific Biosciences) technologies [2, 5, 19, 25, 26, 29]. We have followed the general methodology introduced by mlplasmids [5], that does not rely on databases of known plasmids and chromosomes (Fig. S1). By combining both short and long reads, we create a *hybrid assembly* with UniCycler [28], which typically has a small number of contigs, mostly corresponding to complete circular chromosomes or plasmids. A *short-read assembly* is then constructed from short reads only. The hybrid assembly is used to derive ground truth classification of short-read contigs, as explained in the next paragraph. The short-read assembly graph is then used as an input for our model, both in training and testing scenarios.

**Table 1.** Datasets used in this study. The second column lists the number of samples used for training or testing. Either the SRA project ID or the DOI are listed for data access.

| Species | # samples | Data accession | Reference |
| --- | --- | --- | --- |
| *Bacillus megaterium* | 2 | PRJNA658106 | [26] |
| *Citrobacter freundii* | 88 | PRJNA605147 | [25] |
| *Escherichia coli* | 166 | PRJNA605147 | [25] |
|  |  | PRJNA761884 | [2] |
| *Enterococcus faecium* | 53 | PRJEB28495 | [5] |
|  |  | 10.6084/m9.figshare.7046804 |  |
|  |  | 10.6084/m9.figshare.7047686 |  |
| *Klebsiella oxytoca* | 22 | PRJNA605147 | [25] |
| *Klebsiella pneumoniae* | 45 | PRJNA605147 | [25] |
|  |  | PRJNA761884 | [2] |
| *Staphylococcus pseudintermedius* | 3 | PRJNA521119 | [19] |
| *Vibrio parahaemolyticus* | 3 | PRJEB31923 | [29] |

In hybrid assemblies, the ground truth labels are determined based on the contig length. In particular, all contigs longer than a species-specific threshold (Fig. S2) are labeled as ‘chromosome’, while shorter circular contigs are labeled as ‘plasmid’. The remaining short linear contigs can possibly be a part of an unfinished plasmid or chromosome, and consequently they remain unlabeled. The ground truth labels for short-read assemblies are determined by mapping the contigs to the corresponding hybrid assembly contigs, from which they inherit the labels. The key difference between our pipeline and mlplasmids is that if a contig matches equally well to both chromosomal and plasmidic contigs of the hybrid assembly, it is labeled as ‘ambiguous’. Such contigs are considered as positive examples for both plasmid and chromosome classification tasks. We have observed that without considering such ambiguous matches, the assembly graphs of the short-read assemblies often contained paths with nodes labeled by alternating classes, which is a clearly inconsistent labeling. The use of ambiguous labels allows us to avoid such artefacts. Contigs that matched an unlabeled contig of the hybrid assembly were left unlabeled, and samples that contained more than 5% of unlabeled contigs were discarded from further analysis. The distribution of short read contig labels is shown in Fig. S3. In general, most contigs are labeled as chromosome and less than 1.3% of all contigs are left unlabeled.

Both hybrid and short-read assemblies were created by Unicycler v0.5.0 [28]. Short-read contigs were mapped to the hybrid assembly by minimap2 v2.24 [18] with -c option for accurate alignment.

For training and prediction, all contigs shorter than 100 bp were removed from the short-read assembly graphs and their neighbours were connected by direct edges as part of the feature extraction process. Thus, plASgraph is not predicting the class of contigs shorter than 100 bp. Note that such short contigs often do not have a reliable ground-truth label (often corresponding to SNPs and the like).

## 3 Experimental evaluation

In this section, we evaluate the performance of our plASgraph model and compare it to two recent tools mlplasmids [5] and PlasForest [23] (see Table S1 for overview of experiments). The scripts used for training and evaluation, as well as detailed results are available at https://github.com/cchauve/plASgraph_WABI_2022.

Similarly to plASgraph, mlplasmids is a *de novo* tool, using only sequence-derived features to classify contigs. However, it requires training on each species separately and was designed for contigs longer than 1 kbp. In contrast, PlasForest is a species-agnostic tool, designed to work also on short contigs (*<* 1 kbp), but it is dependent on the comparison of the input sequences to sequence databases. Neither of these tools use the assembly graph information.

Since plASgraph was designed to explicitly handle ambiguous contigs by including separate plasmid and chromosomal classification tasks, we evaluate its prediction accuracy for each of these tasks separately. A contig is predicted as a chromosome if the chromosome score output of the neural network is at least 0.5; and similarly it is predicted as plasmid if the plasmid score is at least 0.5. The true and predicted contig labels then induced the counts of true positives (TP), true negatives (TN), false positives (FP), and false negatives (FN) for each classification task. Each contig was counted as one unit, regardless of its length, and unlabeled contigs were not counted in the evaluation. Ambiguous contigs (those with score *>* 0.5 in both classification tasks) are considered as being labeled both plasmid and chromosome in our accuracy evaluation. As the main accuracy measure, we use the F1 score, which is the harmonic mean of precision (TP/(TP + FP)) and recall (TP/(TP + FN)).

### 3.1 PlASgraph architecture leads to accurate species-specific models

Although our main goal is to produce a single model that can be used across species, we have also trained three species-specific models: *E. faecium* (46 training isolates), *E. coli* (66 training isolates), and *K. pneumoniae* (35 training isolates). The isolates for training were chosen randomly from all available samples except for the *E. faecium*, where we used the same training set as mlplasmids. Out of the training samples, 20% were used as a validation set for the training procedure. The accuracy was evaluated and compared to mlplasmids on held-out testing data sets (*E. faecium* 7 isolates, *E. coli* 100 isolates, *K. pneumoniae* 10 isolates). The development of the model architecture was solely performed on the *E. faecium* data set.

Figure 2 shows that the F1-score of plASgraph is higher than mlplasmids on most samples from *E. faecium* and *K. pneumoniae*. Although mlplasmids outperforms plASgraph on many *E. coli* samples, the overall score for plasmid classification is higher for plASgraph, whereas the performance for the chromosome classification task is comparable (Tab. 2).

**Table 2.**
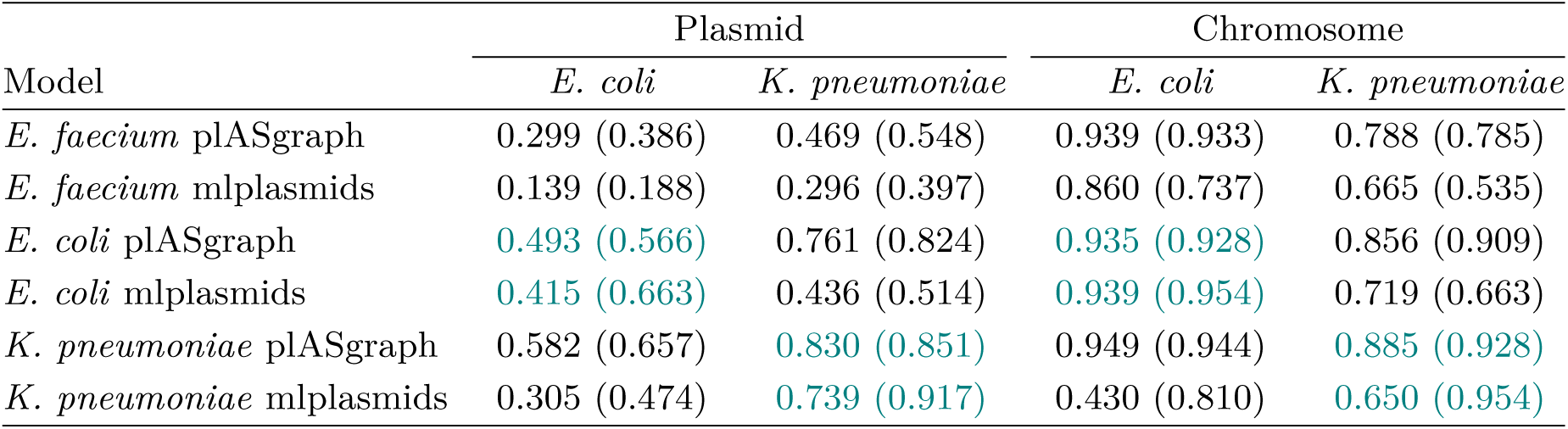
Species-specific and cross-species evaluation. Table shows the average F1-scores across isolates for all analyses of the species-specific models trained (rows) and tested (columns) on a specific combination of species. The cases where the model was trained and tested on the same species are highlighted by teal color. The numbers in parentheses show the results on contigs with length *>* 1 kbp. Plasmid and chromosome classification tasks are evaluated separately. Evaluation on *E. faecium* is not shown, since we have developed the model architecture on this dataset.

**Figure 2.**
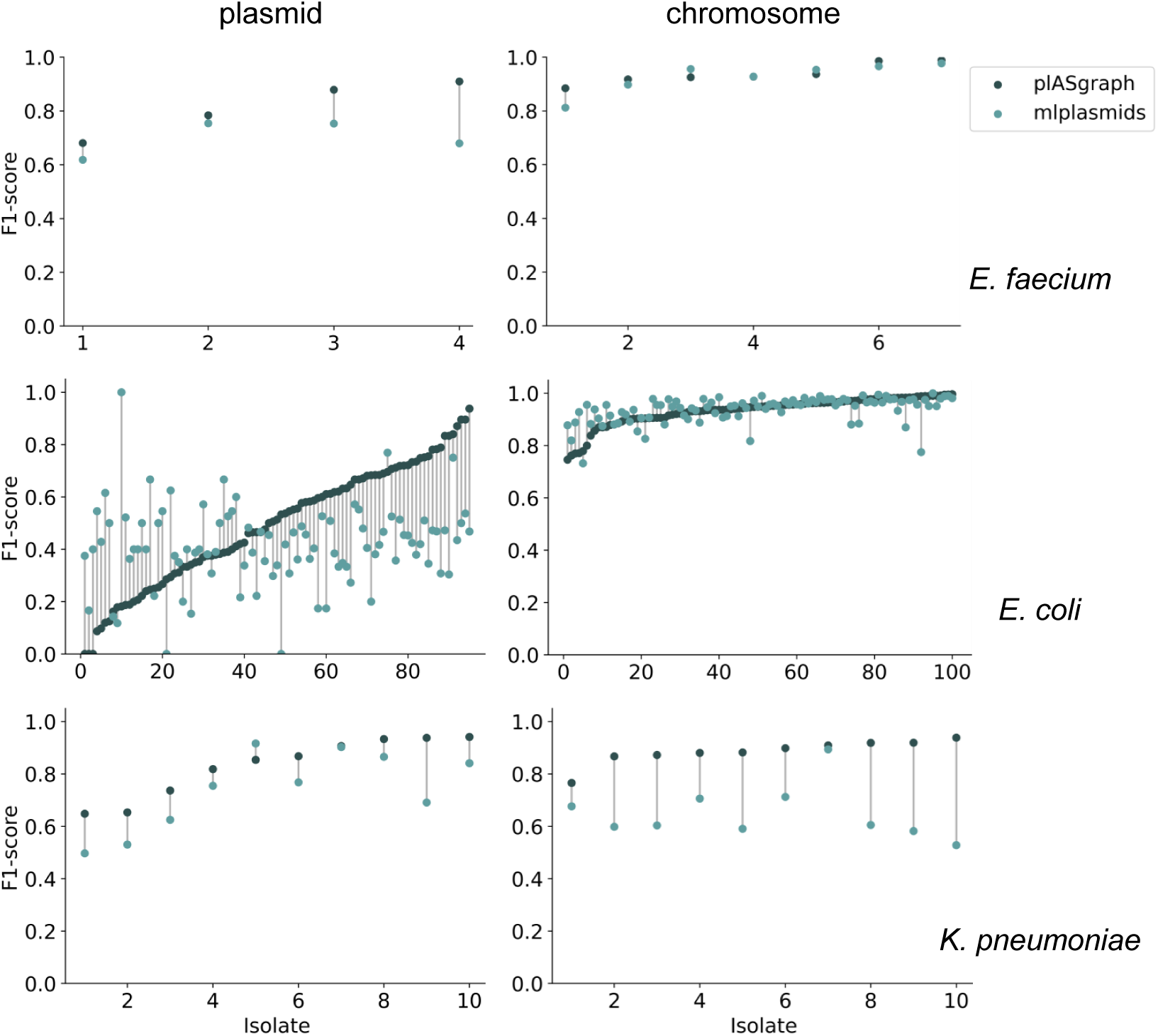
Evaluation of the species-specific plASgraph models in comparison to mlplasmids. Each row compares the F1-score of plASgraph to mlplasmids on individual testing isolates for both plasmid (left) and chromosome (right) classification tasks. The isolates without plasmids are not shown in graphs on the left, as F1-score is then undefined. Rows from top to bottom correspond to *E. faecium, E. coli*, and *K. pneumoniae* data sets respectively. The models are always trained and tested on the same species.

Our results also show that the main advantage gained by plASgraph in the species-specific setting comes from classification of short contigs (100-1000 bp), since considering only contigs above 1 kbp, mlplasmids achieves better accuracy (Tab. 2). This observation can be explained by the *k*-mer frequency feature used by mlplasmids, which provides more detailed information for longer contigs. For contigs shorter than 1 kbp, the *k*-mer frequency feature may only contain counts for a few distinct *k*-mers. This can lead to a relatively high number of zero values in the feature vector which does not allow the model to classify the respective contig accurately. In contrast, plASgraph uses only a single *k*-mer related feature, which is supplemented by information from graph neighbourhood; this combination likely helps to classify short contigs more accurately. Figure 3 demonstrates that indeed, the addition of GCLs significantly increases the prediction accuracy compared to a simpler model with the same input features.

**Figure 3.**
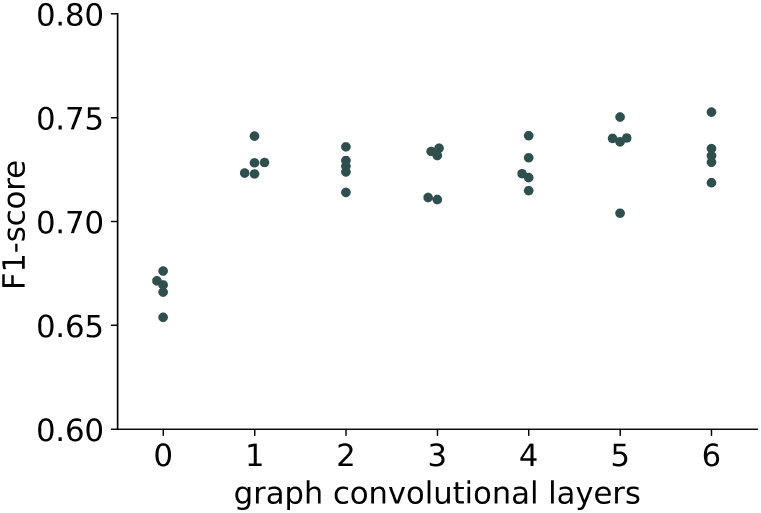
Impact of the number of layers on the model performance. Different numbers of graph convolutional layers (GCLs) were tested for the *E. faecium* model. Each architecture was trained five times using random seeds and the F1-score was calculated on the 20% split validation set. The largest improvement in performance is visible with introduction of the first GCL; a slight additional increase in accuracy is observed for five and six GCLs.

### 3.2 Species-specific plASgraph models generalize well to other species

Table 2 also shows that in cross-species application of species-specific models, plASgraph has a distinct advantage compared to mlplasmids. For example, the plASgraph model trained on *E. coli* achieves plasmid F1-score 0.76 on *K. pneumoniae*, which is only slightly lower than F1-score 0.83 achieved by the model trained on the same species. In the case of mlplasmids, we see a significant decrease of the F1-score, from 0.74 on the same species to 0.44 across species. Similar trends persist when we restrict evaluation to contigs above 1 kbp. Note the *E. faecium* models are less accurate on *E. coli* and *K. pneumoniae* testing sets due to the large phylogenetic distance, but plASgraph still performs significantly better than mlplasmids. The distribution of per sample F1-scores is shown in Fig. S4.

We attribute this better cross-species generalization of plASgraph to our use of input features that are relative to the sample-wide statistics, and thus are less species dependent and are able to use the overall context provided by the whole assembly. In contrast, mlplasmids directly uses *k*-mer frequencies, which are highly specific to individual species.

### 3.3 Species-agnostic plASgraph model

One of the goals of the plASgraph model was to create a tool that could be applied to newly identified species for which no information is available in sequence databases and no training sets assembled with long reads are readily available. To this end, we have trained a species-agnostic model on a mixed training set from *E. faecium, E. coli*, and *K. pneumoniae*, using 20 isolates from each as a part of the training set. Again, 20% of the training set was withheld for validation during training. We evaluated the performance of the species-agnostic model on five species, different from those included in the training set. All available isolates for the five species (*B. megaterium* (2), *C. freundii* (88), *K. oxytoca* (22), *S. pseudintermedius* (3) and *V. parahaemolyticus* (3)) were used for testing. Since mlplasmids is not suitable for this application, we compared the plASgraph model with the recent PlasForest model [23] which has been designed for cross-species use.

Figure 4 shows that plASgraph has better average plasmid accuracy than PlasForest on four out of five species (see also Fig. S5 for the distribution of F1-scores for individual isolates). For the *S. pseudintermedius* data set, which contained only a single plasmid, both tools failed to predict that plasmid correctly. Both tools have high accuracy in predicting chromosomal contigs, with plASgraph being more accurate on *C. freundii, K. oxytoca*, and *B. megaterium*, but performing slightly worse for the remaining two species. The performance for ambiguous contigs is shown in Figure S6. Figure S7 shows scatter plot of all *C. freundii* contigs based on their chromosome and plasmid score and their true label. It shows that overall plASgraph classifies a large majority contigs accurately.

**Figure 4.**
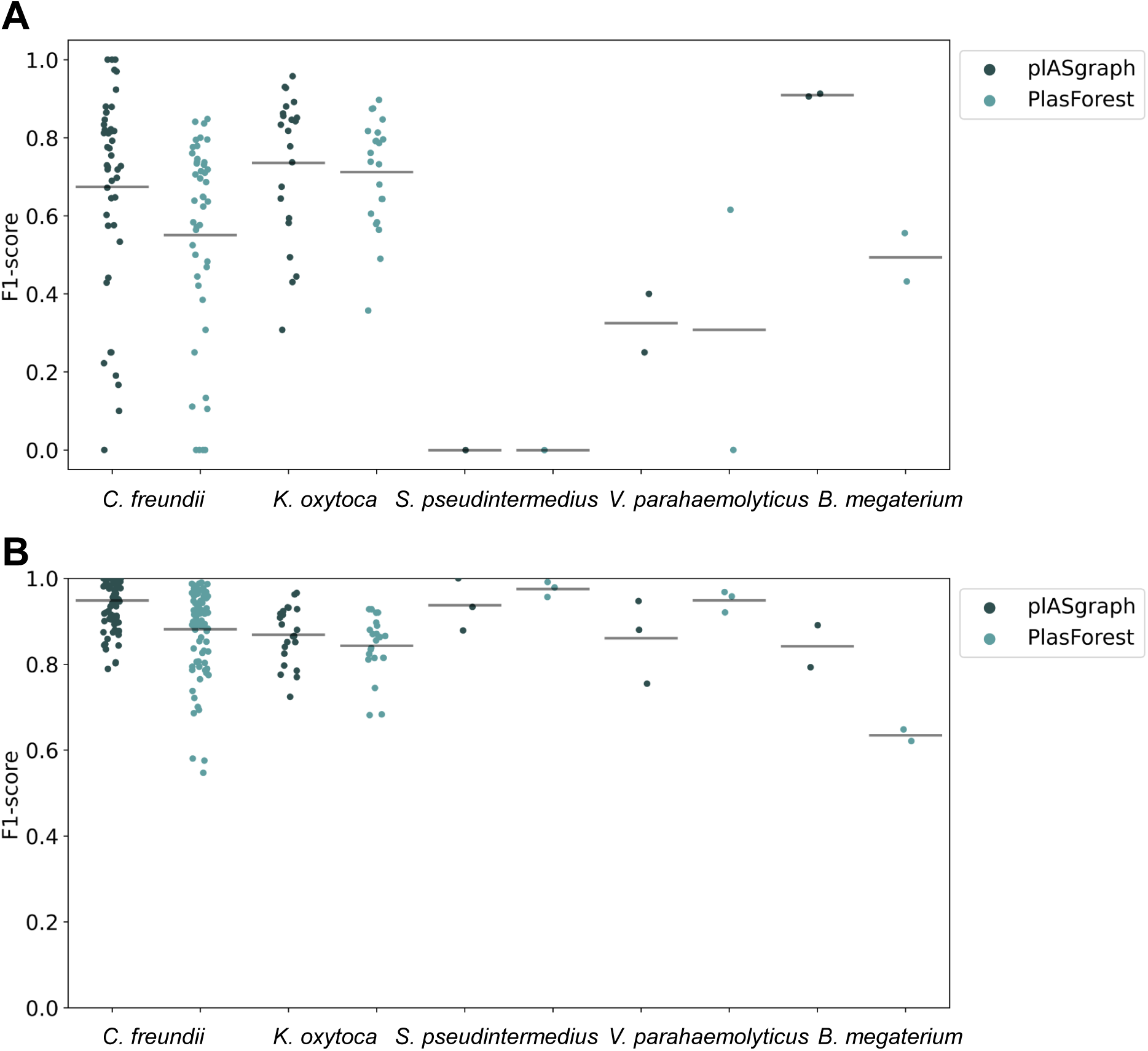
Evaluation of the species-agnostic plASgraph model and comparison to PlasForest. A: plasmid classification, B: chromosome classification. Horizontal lines represent the average F1-score of all isolates used for testing.

PlasForest uses features derived from querying input sequences against a reference database, which is not used in our predictions. The higher average F1-score of plASgraph therefore suggests that the contribution of information present in the assembly graph combined with relative features of the contigs exceeds the contribution from homology search. Moreover, independence from sequence databases makes our tool more suitable for application to completely novel species.

PlASgraph not only provides a score for plasmidic and chromosomal contigs but also outputs a visualization of an assembly graph labeled according to the predictions. Figure 5 shows parts of the assembly graph for *C. freundii* isolate SAMN15148288 with nodes colored according to the ground truth and both plASgraph and PlasForest predictions. The ground truth supports our initial reasoning to incorporate the information provided in the assembly graph, as linked contigs are more likely to belong to the same class. While both tools make some incorrect predictions, visualization clearly shows several isolated chromosome predictions among plasmid contigs and vice versa in the PlasForest prediction, whereas plASgraph has only one such isolated false positive. In general, plASgraph predictions are more consistent with the assembly graph topology.

**Figure 5.**
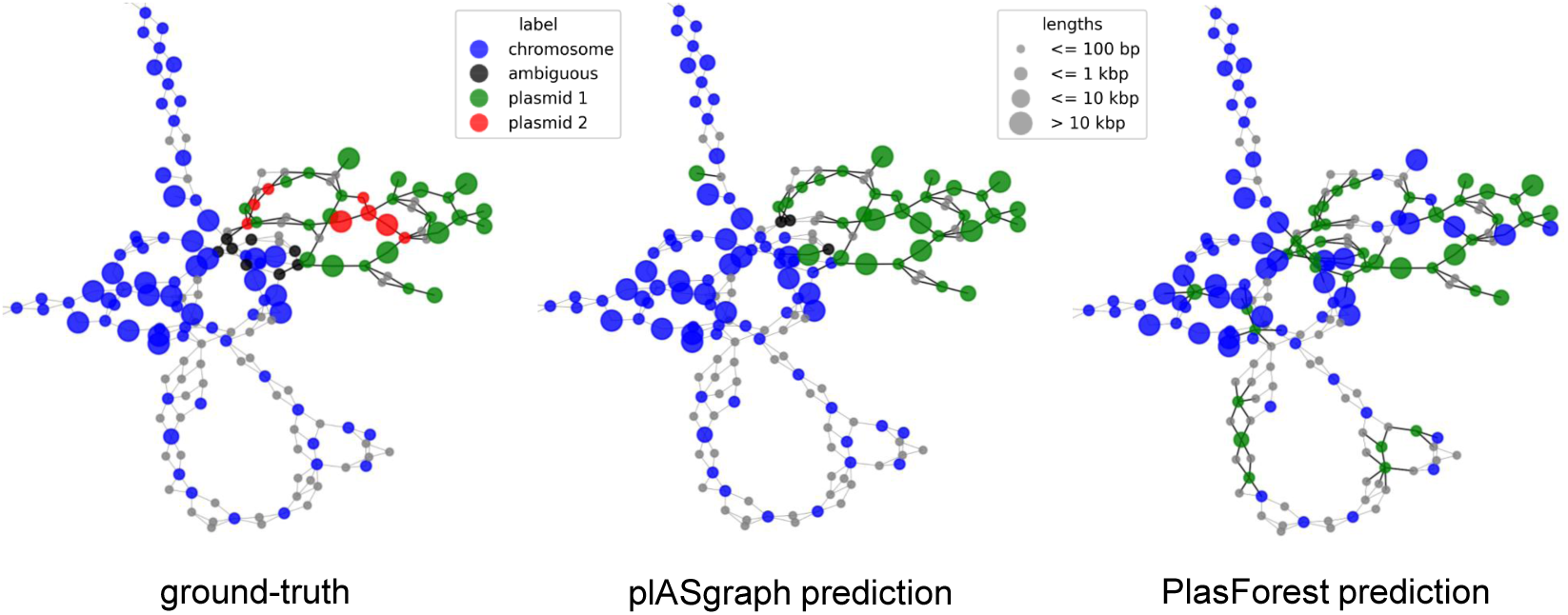
Contig classification in the context of the assembly graph of *C.freundii* isolate SAMN15148288. Chromosomal contigs are colored in blue and ambiguous contigs are colored in black. Left: The ground-truth, including two different plasmids (green and red). Middle: plASgraph predictions. Right: PlasForest predictions. Note that the classification tasks do not include binning of contig plasmids, thus all predicted plasmid contigs are color in green. The assembly graph extends to the upper left as a loop of chromosomal contigs alternating with unlabelled SNPs, which is not shown.

## 4 Discussion

PlASgraph is a GNN that can be used to identify plasmidic, chromosomal, and ambiguous contigs directly from a bacterial assembly graph. Our tool is easy to use, only requires a short-read assembly graph file as an input, and outperforms other state-of-the-art methods.

PlASgraph is not dependent on specific species and can therefore be used also for newly sequenced bacteria for which no closed genome sequence is available yet. Our species-agnostic plASgraph model outperforms recently published, database-dependent PlasForest [23]approach when compared across different bacterial species. *De novo* classification (database independence) allows more accurate identification of previously unknown plasmids and chromosomes which can be critical for diverse One-Health epidemiologic surveillance.

However, when desired, plASgraph can also be trained for a particular species. Our species-specific models are more accurate when compared to mlplasmids [5], although mlplasmids performs better than plASgraph on longer contigs above 1 kbp. We hypothesize that mlplasmids is able to learn to recognize chromosome contigs of a particular species through their pentamer distributions at the cost of cross-species generalizability. Furthermore, this approach is unreliable for classifying contigs of lengths in the range of 100-1000 bp. Accurate classification of shorter contigs by plASgraph may enable identification of more complete plasmids from incomplete assemblies and has a potential to facilitate novel plasmid discovery.

Another novel feature of plASgraph is the separation of plasmid and chromosome classification tasks, recognizing that some contigs are ambiguous, being parts of both types of molecules. These ambiguous contigs are an interesting subject for further study by themselves; our preliminary analysis of ambiguous contigs in our datasets suggests that the majority of them are related to transposons and phages. These mobile elements can integrate into both plasmids and chromosomes within the cell.

The simplicity of the architecture of the plASgraph model makes it amenable to extensions. For example, the use of additional information about plasmids, such as the presence of plasmid-specific genes in a contig, could allow further increase in classification accuracy as this additional information would propagate to nearby nodes thanks to the GNN architecture. In addition, it will be interesting to investigate how plASgraph could be adapted for accurate plasmid identification in metagenomic datasets, like wastewater samples, which play an increasingly crucial role in monitoring antibiotic resistance [13].

## Acknowledgements

This research was enabled in part by support provided by Compute Canada (www.computecanada.ca). We thank Aniket Mane for introducing the idea of using GNN with assembly graphs. Most of the work was conducted during a visit of JS, KS, BB and TV to Simon Fraser University enabled by the PANGAIA EU project.

## 5 Supplementary figures and tables

**Figure S1.**
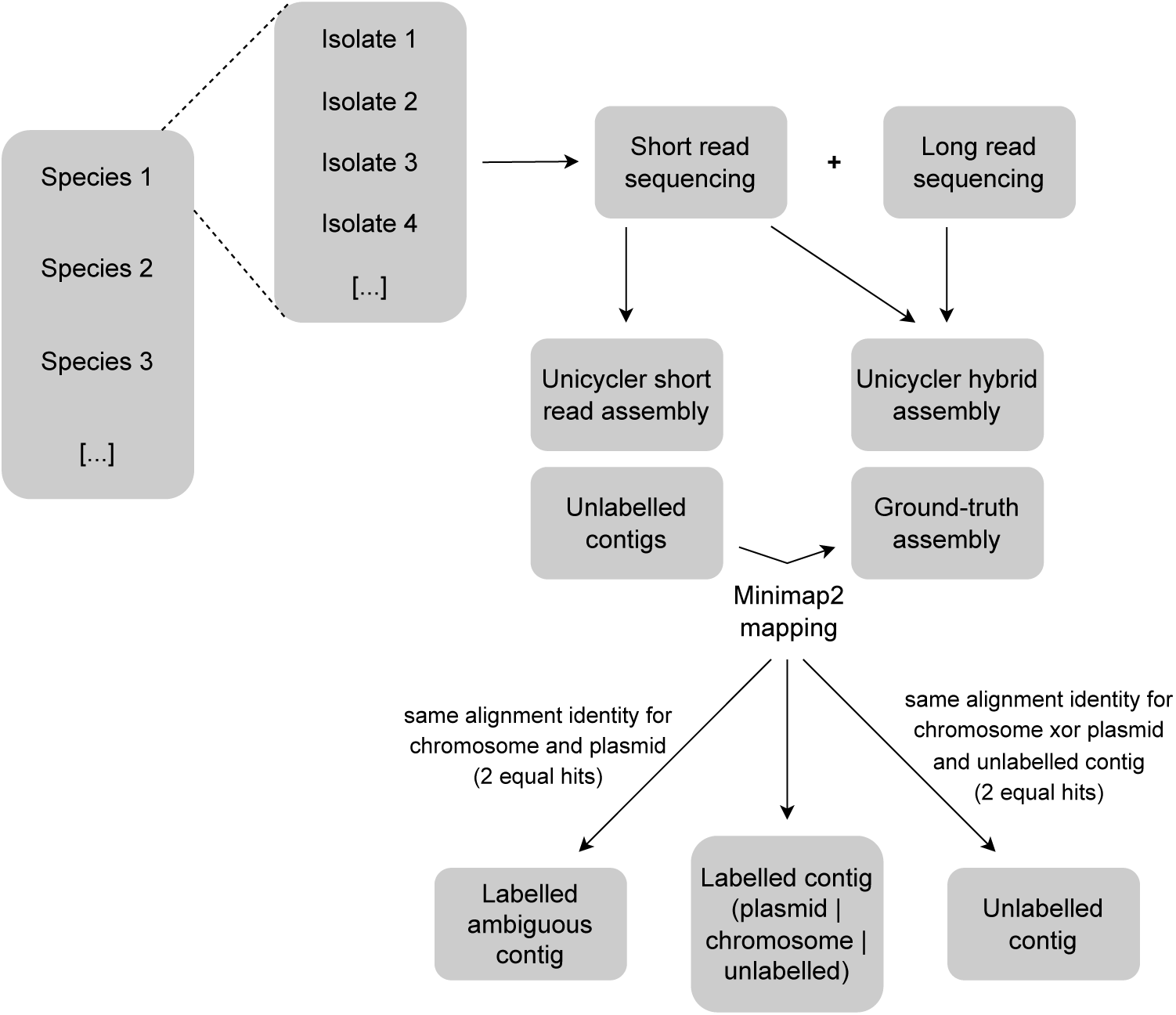
Labeling workflow. A short-read and hybrid assemblies are generated for each isolate. Unlabeled contigs of the short-read assembly are then mapped against the ground-truth hybrid assembly using minimap2 [18]. In case of a unique best alignment, the short-read contig is labeled according to the matching hybrid contig. If two equally good hits are identified to a chromosome and a plasmid hybrid assembly contig, the short-reads contig is labeled as ‘ambiguous’. If two equally good hits are identified to a chromosome or a plasmid contig and to an unlabeled contig, the contig is ‘unlabeled ‘.

**Figure S2.**
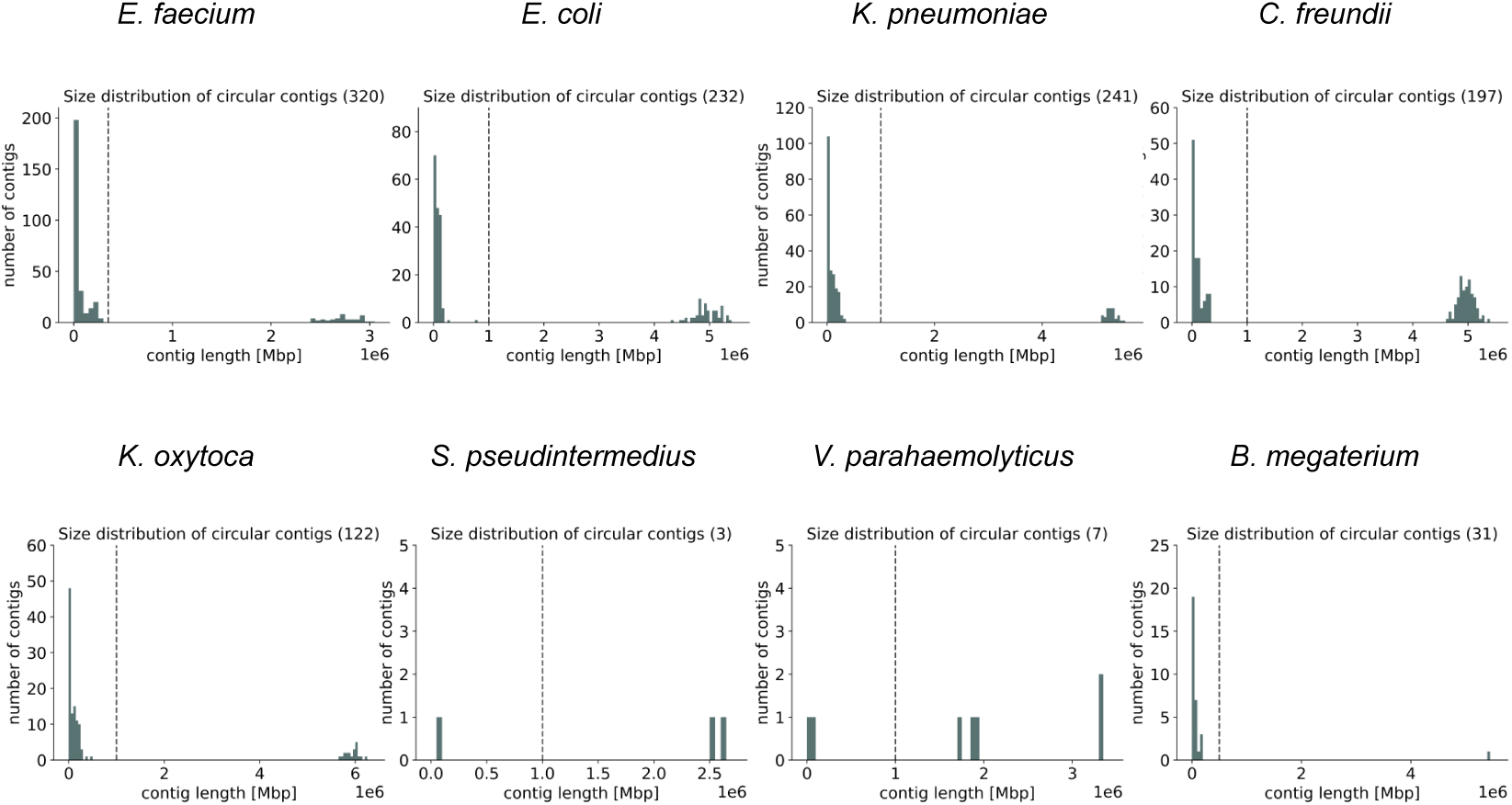
Circular contig size distribution in hybrid assemblies. Above each plot, the total number of circular contigs is shown in parentheses. A vertical line marks the species-specific threshold in each plot; circular contigs shorter than the threshold are considered plasmidic, whereas longer contigs are considered chromosomal.

**Figure S3.**
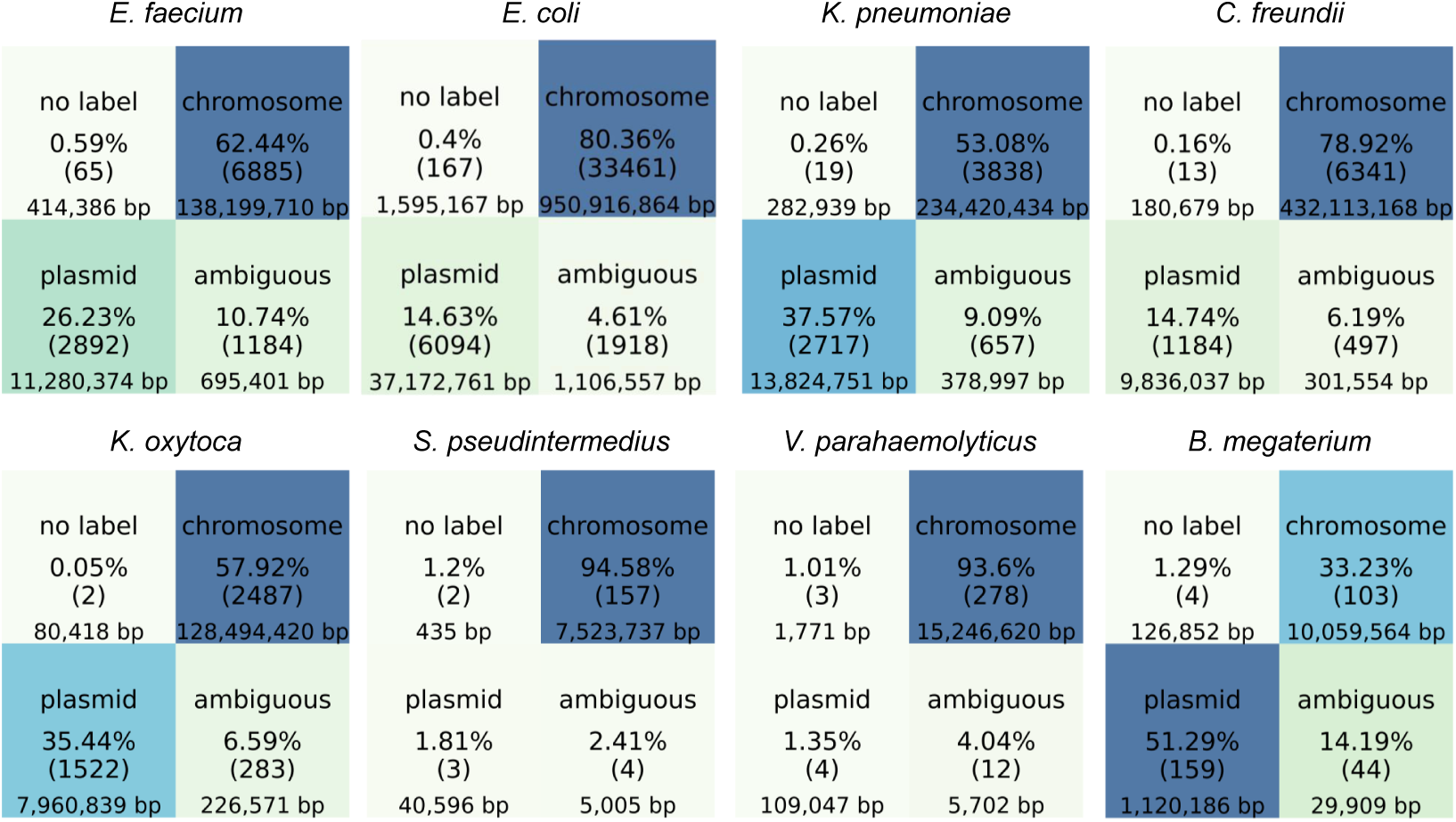
Distribution of short-read contig labels. Darker colors represent a higher percentage of the respective label in each dataset. Each small square shows the percentage, the absolute number, as well as the cumulative length of the short-read contigs with the respective label.

**Table S1.**
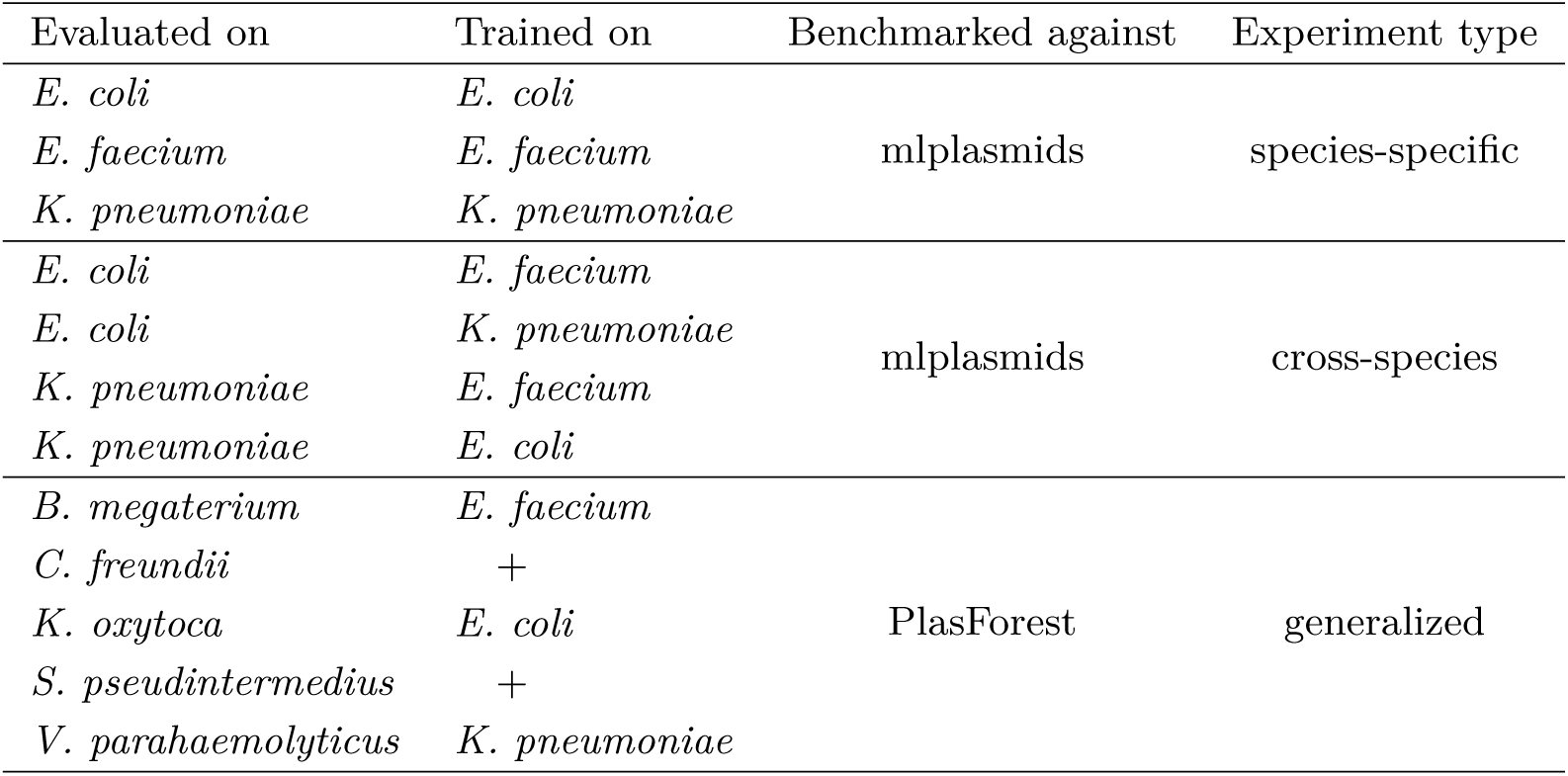
Overview of the performed comparative analyses.

**Figure S4.**
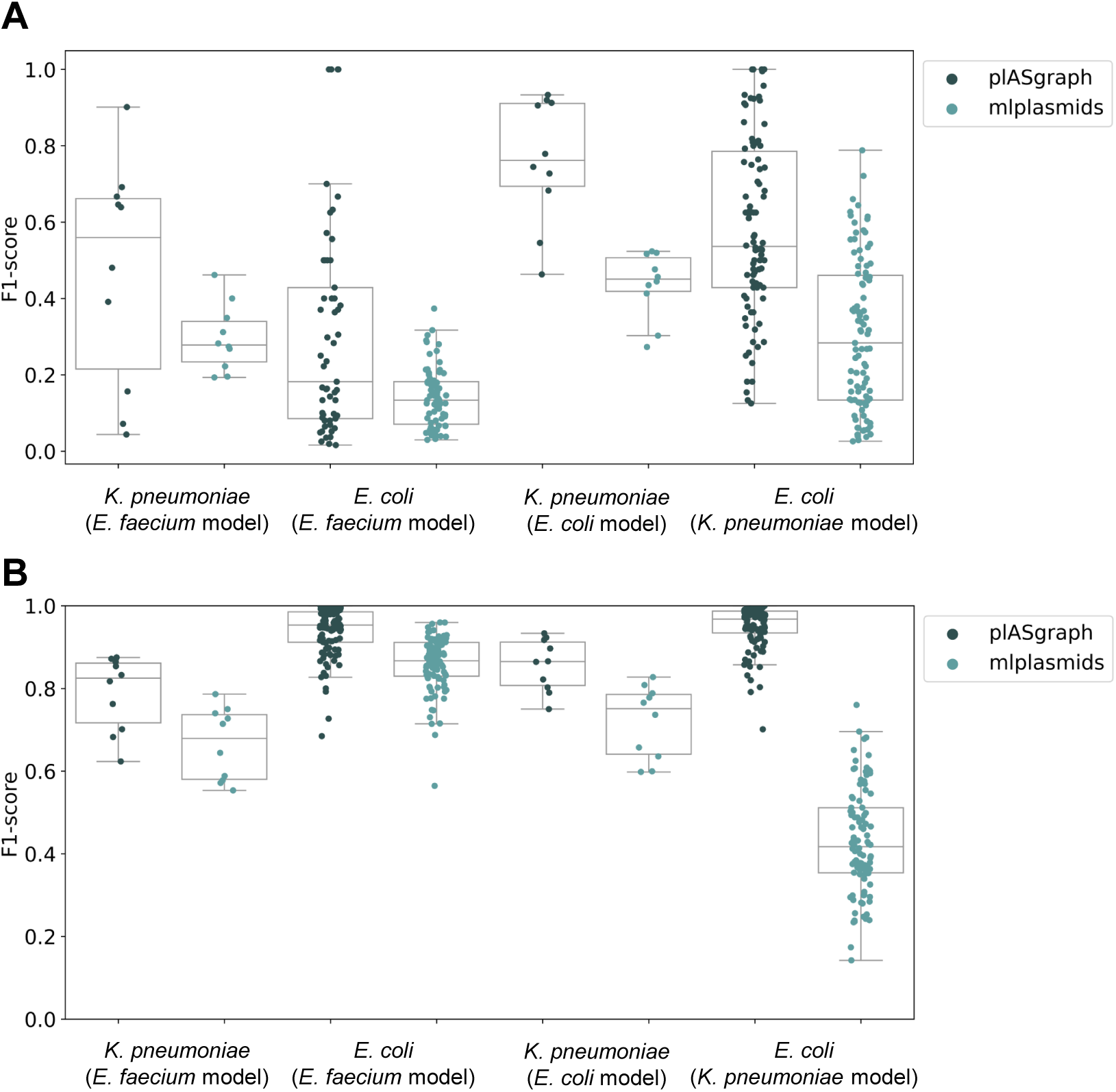
Cross-species evaluation of the species-specific plASgraph models in comparison to mlplasmids. A: Plasmid classification. B: Chromosome classification.

**Figure S5.**
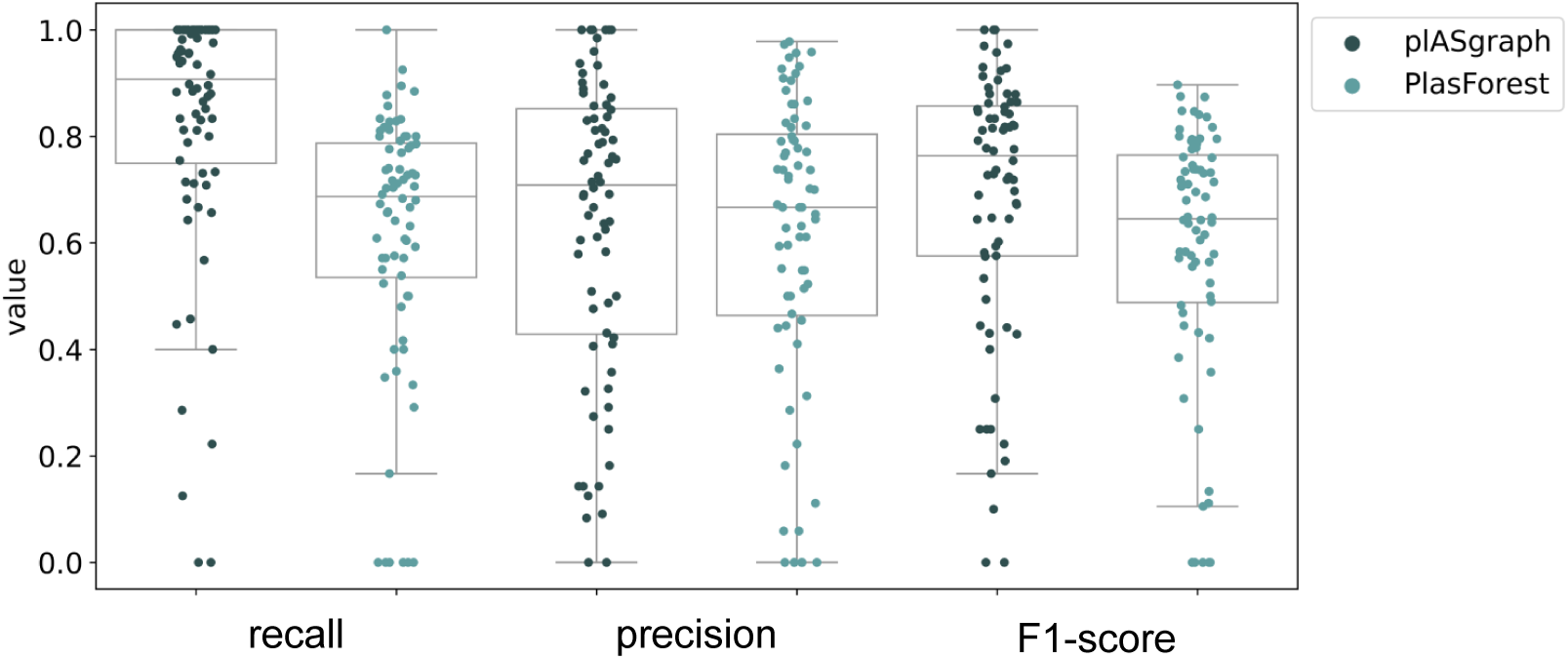
Accuracy of plasmid classification for species-agnostic plASgraph model in comparison to PlasForest. Each data point represents one isolate; the plot combines isolates from all species used in Fig. 4.

**Figure S6.**
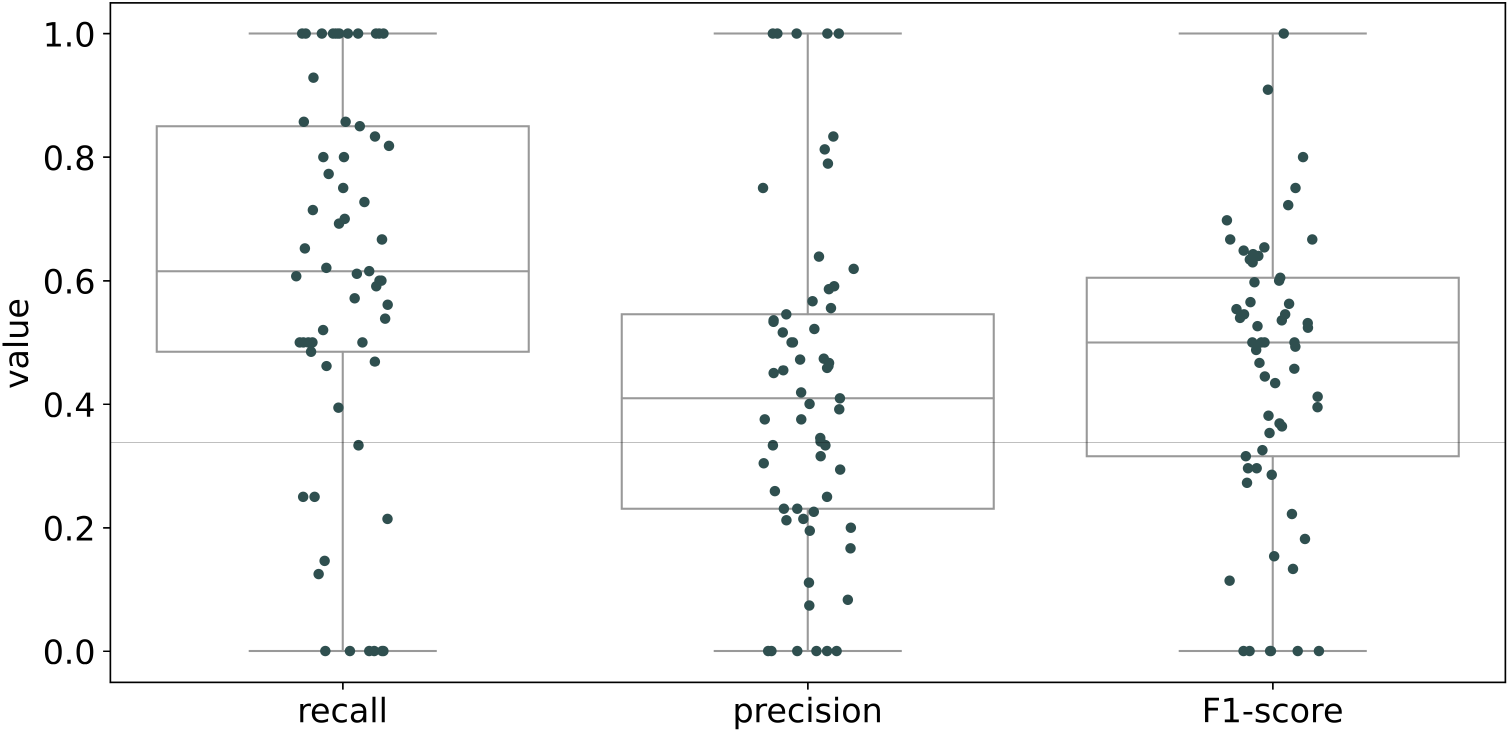
Accuracy of ambiguous contig classification for species-agnostic plASgraph model. Each data point represents one isolate; the plot combines isolates from all species used in Fig. 4. The isolates with no ambiguous contigs are not shown.

**Figure S7.**
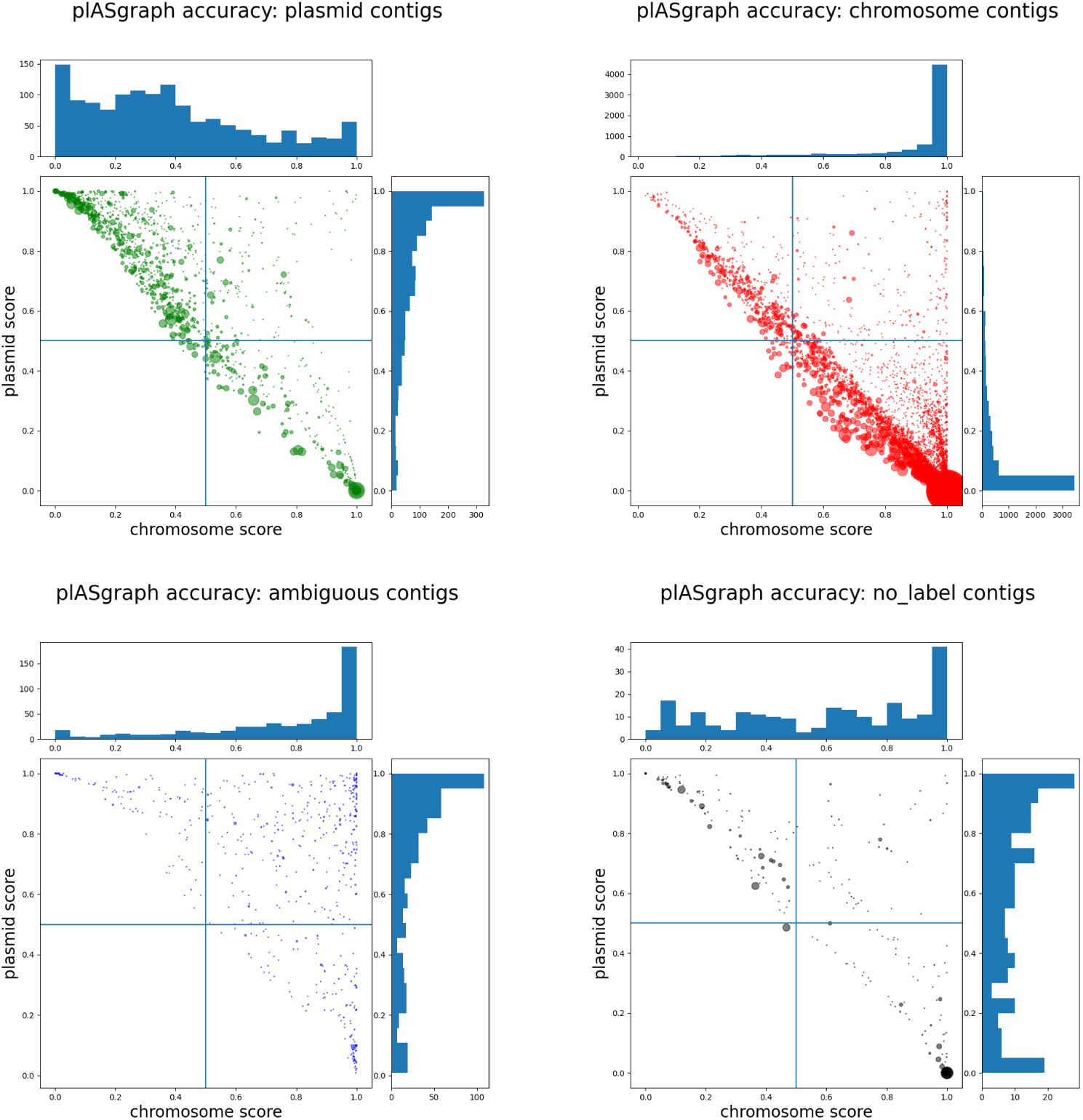
Results on the 96 *C. freundii* samples (9118 contigs). Each dot represents a contig, its radius being proportional to the contig length. Contigs are split in four panels according to their ground truth label. Each contig is shown with coordinates being its chromosome score (x-axis) and its plasmid score (y-axis).

